# Insights on the impact of mitochondrial organisation on bioenergetics in high-resolution computational models of cardiac cell architecture

**DOI:** 10.1101/327254

**Authors:** Shouryadipta Ghosh, Kenneth Tran, Lea M. D. Delbridge, Anthony J. R. Hickey, Eric Hanssen, Edmund J. Crampin, Vijay Rajagopal

## Abstract

Recent electron microscopy data have revealed that cardiac mitochondria are not arranged in crystalline columns, but are organised with several mitochondria aggregated into columns of varying sizes often spanning the cell cross-section. This raises the question - how does the mitochondrial arrangement affect the metabolite distributions within cardiomyocytes and their impact on force dynamics? Here we employed finite element modelling of cardiac bioenergetics, using computational meshes derived from electron microscope images, to address this question. Our results indicate that heterogeneous mitochondrial distributions can lead to significant spatial variation across the cell in concentrations of inorganic phosphate, creatine (Cr) and creatine phosphate (PCr). However, our model predicts that sufficient activity of the creatine kinase (CK) system, coupled with rapid diffusion of Cr and PCr, maintains near uniform ATP and ADP ratios across the cell cross sections. This homogenous distribution of ATP and ADP should also evenly distribute force production and twitch duration with contraction. These results suggest that the PCr shuttle, and associated enzymatic reactions, act to maintain uniform force dynamics in the cell despite the heterogeneous mitochondrial organization. However, our model also predicts that under hypoxia - activity of mitochondrial CK enzyme and diffusion of high-energy phosphate compounds may be insufficient to sustain uniform ATP/ADP distribution and hence force generation.

## Author Summary

Mammalian cardiomyocytes contain a high volume of mitochondria, which maintains the continuous and bulk supply of ATP to sustain normal heart function. Previously, cardiac mitochondria were understood to be distributed in a regular, crystalline pattern, which facilitated a steady supply of ATP at different workloads. Using electron microscopy images of cell cross sections, we recently found that they are not regularly distributed inside cardiomyocytes. We created new spatially accurate computational models of cardiac cell bioenergetics and tested whether this heterogeneous distribution of mitochondria causes non-uniform energy supply and contractile force production in the cardiomyocyte. We found that ATP and ADP concentrations remain uniform throughout the cell because of the activity of creatine kinase (CK) enzymes that convert ATP produced in the mitochondria into creatine phosphate, which rapidly diffuses to the myofibril region where it can be converted back to ATP for the contraction cycle in a timely manner. This mechanism is called the phosphocreatine shuttle (PCr shuttle). The PCr shuttle also ensures that different areas of the cell produce the same amount of force regardless of the mitochondrial distribution. However, our model also shows that when the cellular oxygen supply is limited - as can be the case in conditions such as heart failure - the PCr shuttle cannot maintain uniform ATP and ADP concentrations across the cell. This causes a non-uniform acto-myosin force distribution and non-uniform twitch duration across the cell cross section. Our study suggests that mechanisms other than the PCr shuttle may be necessary to maintain uniform supply of ATP in a hypoxic environment.

## Introduction

Cardiomyocytes require a ready supply of adenosine triphosphate (ATP) in order to generate the contractions that cause the heartbeat. ATP demands at hydrolysis sites within myofibrils can vary five-fold between resting and active states of heart [1]. As a result, cardiomyocytes are densely packed with mitochondria, which traverse in between columns of contractile proteins called myofibrils. Previous studies suggested that cardiac mitochondria are arranged in a ‘crystal like’ regular pattern with very small deviation in the distances between neighbouring parallel strands of mitochondria [2, 3]. It was also predicted that this highly ordered mitochondrial arrangement further leads to the formation of functional units called Intracellular energetic units (ICEU) [3–5]. Within the ICEUs, neighbouring mitochondria and myofibrils were proposed to interact via energy transfer mechanisms including phosphocreatine (PCr) shuttle facilitated transport of ATP/ADP, direct diffusion of ATP/ADP as well as adenylate kinase reactions. It was proposed that there was no significant exchange of ATP/ADP with the larger cytosolic pool of high-energy phosphate compounds (phosphagens) outside the ICEUs.

However, these previous studies were based on confocal image data, which did not provide an accurate description of the mitochondrial arrangement. We recently analysed [6] 2D electron microscopy (EM) images of cross-sections covering the entire diameter of the cell from Sprague Dawley (SD) rat hearts. The analysis revealed that cardiac myocytes have a non-crystalline distribution of mitochondria across the cross sections. Individual mitochondria varied in shape, size and exhibited variations in clustering between the myofibrillar bundles. We also found that there is a statistically significant difference in the size of the mitochondrial clusters between control and STZ induced type1-diabetic hearts [6]. Using a compartmental model of mitochondrial oxidative phosphorylation, we proposed that these changes in mitochondria morphology and distribution could moderately enhance energy supply in diabetic cardiomyopathy.

Other recent EM studies [7, 8] also confirm our observations about the heterogeneous distribution of mitochondria in cardiomyocytes. Considering the previously proposed hypothesis of ICEUs, this observed mitochondrial heterogeneity could give rise to a non-uniform distribution of ATP/ADP which can have a significant impact on the contractile performance of the cell. Furthermore, we [6] and others [8, 9] have reported changes to this heterogeneous distribution of mitochondria as a consequence of changes to mitochondrial substrate selection and metabolism in disease conditions. Thus it is important to understand how the cell maintains a uniform distribution of ADP, ATP and acto-myosin force production with a heterogeneous mitochondrial distribution.

In this study, we build on our earlier 2D stereological and compartmental model simulation analyses in Jarosz et al [6] and investigate the effect of the mitochondrial distribution displayed in these high-resolution images on the spatial distribution of phosphagens and contractile force dynamics across the cross-section of the cell. We have studied the diffusion of phosphagens within the cell cross section using new, spatially detailed finite element models. Although finite element (FE) based computational models have been used to study the energy landscape in cardiac myocytes previously [10–13], these models defined idealised geometric domains (crystalline mitochondria or rectangular grid-like arrangements) and did not account for the mitochondrial clustering and non-uniform distribution that we observe in our data (Fig 1.). The simulations and analyses in our study were conducted using three cross-section images that were selected from different positions within a 3D serial block-face scanning electron microscopy (SBF-SEM) dataset of a single control cardiomyocyte (Fig 1). This allowed us to compare between simulation results from different regions of the cell and to investigate the various structural factors that influence metabolic landscape of the cell.

**Fig 1.**
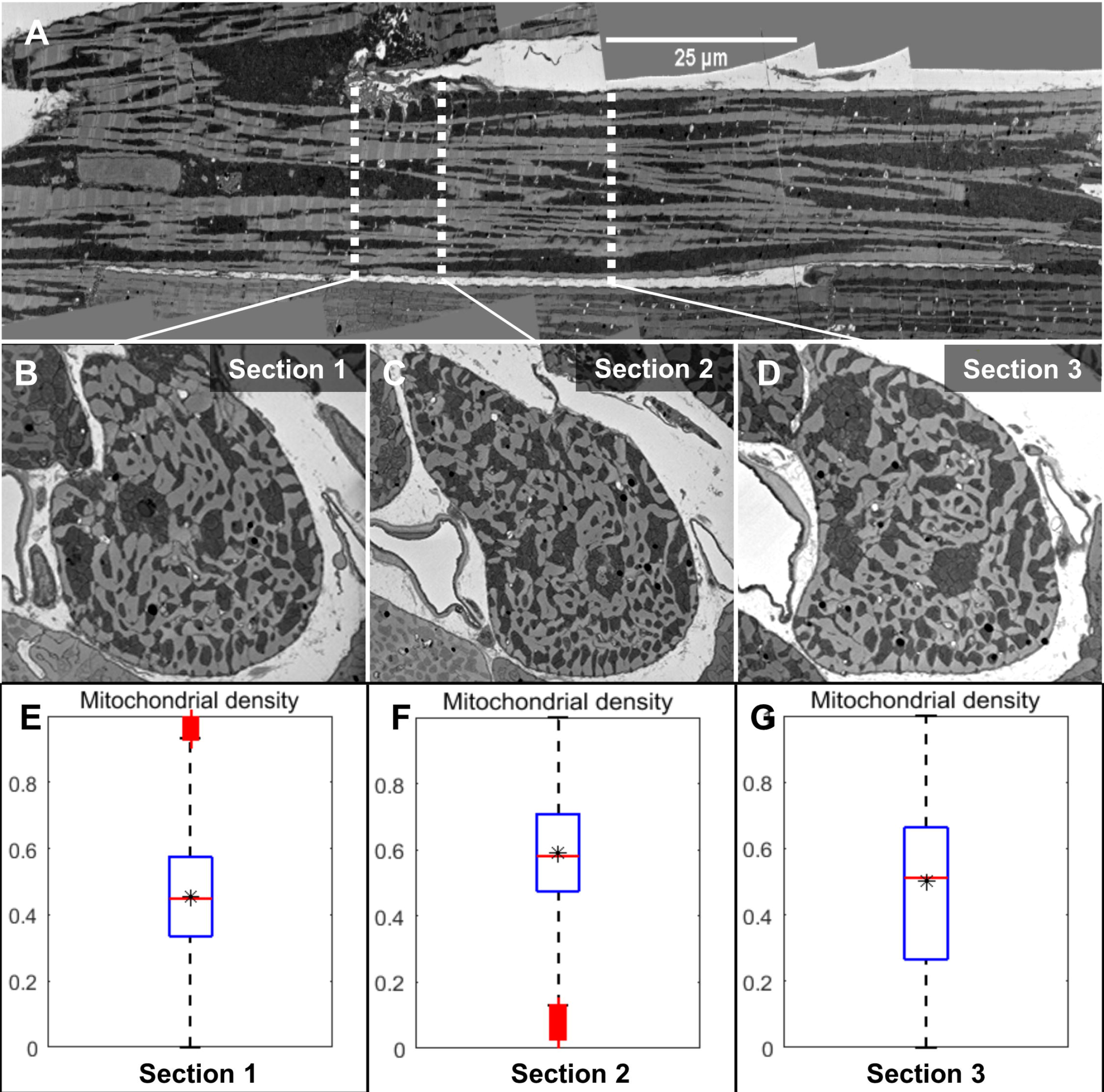
EM image of a single cardiac myocyte and analysis of mitochondrial distribution in different cross-sections. (A): A Longitudinal section from the SBF-SEM images of a single cardiomyocyte with mitochondria in dark regions and myofibril/nucleus in light regions. (B-D): Cross sections (1-3) corresponding to the location of three white dotted lines in the longitudinal section. Section 1 is located near a branching point, and section 2 is located close to a nucleus, however none of the sections contain a nucleus (E-G): Statistical box plot representing the distribution of mitochondrial density at various points in the cross-section 1 -3. The variation in mitochondrial density distribution is highest in section 3 and lowest in section 1. The stars represent the total mitochondrial area fraction present in each cross section.

When investigating how the heterogeneous distribution of mitochondria affects the distribution of ATP, we must also account for the roles that the phosphocreatine shuttle mechanism and oxygen (O_2_) supply play in maintaining energy supply within the cell. A uniform supply of O_2_ would ensure that each mitochondrion in the cell cross section can accomplish the final stage of oxidative phosphorylation to produce ATP. The phosphocreatine shuttle enables rapid transport of inorganic phosphate from the vicinity of mitochondria to the vicinity of myofibrils, where it is used for muscle contraction in the form of ATP. Models such as [10–13], have only studied the role that phosphocreatine shuttle plays in the context of a uniform or crystalline distribution of mitochondria. To the authors’ knowledge no spatially realistic computational model has yet examined how oxygen supply and distribution across the cell cross section (or lack thereof in hypoxic conditions) might also interact with the phosphocreatine shuttle mechanism to facilitate uniform supply of ATP. Specifically, we set out to determine whether the phosphocreatine shuttle mechanism is sufficient to maintain uniform ATP supply across the cross section in normoxic and hypoxic conditions at moderate and high workloads in a heterogeneously distributed mitochondrial environment.

We report several new and important findings based on the computational studies that we have conducted in this study: (i) The heterogeneous mitochondrial distribution can lead to large gradients in the concentration of inorganic phosphate (Pi), creatine phosphate (PCr) and creatine (Cr) at high workloads when exposed to normal oxygen supply (normoxia) from the capillaries. These large concentration gradients exist over the entire cross-section of the cell between areas with high and low localised mitochondrial density. However, the gradients in ATP and ADP are negligible due to the rapid diffusion of PCr and Cr coupled with the activity of CK enzymes; (ii) regional variation in intracellular oxygen supply can impact the metabolite distribution under hypoxic conditions; (iii) and finally, by computing the distribution of peak force and twitch duration using our simulated metabolite distributions as input, we show that the peak force distribution and twitch duration is insensitive to a heterogeneous distribution of phosphagens under normoxic conditions. However, the twitch duration can vary by as much as 30 ms between different parts of the cell under hypoxic conditions.

The following sections present the formulation of the finite element model simulations and the subsequent results that point towards the above-mentioned findings. In the results section, we first analysed the localised density of mitochondria within the three cross-sections that were used for this study. We have also provided details of the model formulation along with validation of the model with experimental results. Following this, we have presented the results from six different sets of simulations based on the three cross-sections under normoxic and hypoxic conditions. We discuss the physiological significance of the presented results, limitations of the current analysis and future directions. Finally, the mathematical formulation of the FE model used in these simulations is described in detail in the material and methods section.

## Results

### Quantification of mitochondrial distribution in single cell cross-sections

We collected serial block face scanning electron microscopy (SBF-SEM) images of one complete cell [14, 15]. Fig 1A presents a longitudinal section from the 3D dataset, while Figs 1B, 1C and 1D show the cross sections corresponding to the positions marked by the white lines on Fig. 1A. The first cross-section in Fig 1B (section 1) was located near a branching point of the cell. On the other hand, section 2 is located in the proximity of a nucleus, although none of the three cross-sections contain any portions of the nucleus. As evident from these images, mitochondria were highly clustered in both lateral and longitudinal sections. In order to quantify the distribution of mitochondria in the cross sections, we manually segmented the cell membrane and mitochondrial cluster boundaries [16] in the three cross sectional images (S1A Fig). Following this, we calculated the localized area density of mitochondria (simply referred as mitochondrial density here on) at different points in the images, which is defined as the normalized mitochondrial area (Area_M_) present inside an 1.6 μm × 1.6 μm square local window (Area_T_) centred at a given point (S1B Fig). We also tested the sensitivity of this measurement to the size of the square window used in the analysis. The difference between the first and third quantile of the mitochondrial density distribution was maximum when a small local window size (<1 μm) was used (S1C Fig) and decreasing as the window size increased. Increasing the window size also leads to a decrease in the difference between median of the mitochondrial density distribution and the fractional area of the total mitochondria present in the cross section. However, increasing the window size beyond 1.6 μm did not significantly change the median of the distribution, and it also resulted in a large number of outliers. Therefore, a window size of 1.6 μm was used for our analyses (Fig 1E-G).

We found a significant variation in the mitochondrial density distribution of each cross section (Fig 1E-G). The first and third quantile of the distribution differs by a margin of 20% - 40% in these images. The margin of this difference is highest in section 3 and lowest in section 1. The star marks inside the boxplots represent the fractional area of total mitochondria present in these sections. Section 2 has the highest area fraction of mitochondria (59%), followed by section 3 (50%) and section 1 (45%).

### A high resolution two-dimensional spatial model of cardiac bioenergetics and its validation with experimental results

These descriptions of mitochondrial distribution were incorporated into a 2D finite element (FE) model of cardiac bioenergetics. We first assumed that the distribution of phosphagens within the cardiomyocyte cross sections will be similar across the length of several sarcomeres. We further assumed that individual mitochondria that cluster together share a common inner membrane space (IMS) (Fig 2A). We made this assumption based on a previous EM imaging study by Picard et al. [17], who reported that adjacent mitochondrial outer membranes in cardiac myocytes are connected through electron dense inner-membrane junctions (IMJ). Picard et al. also found a close alignment between the cristae belonging to adjacent mitochondria along the IMJ.

**Fig 2.**
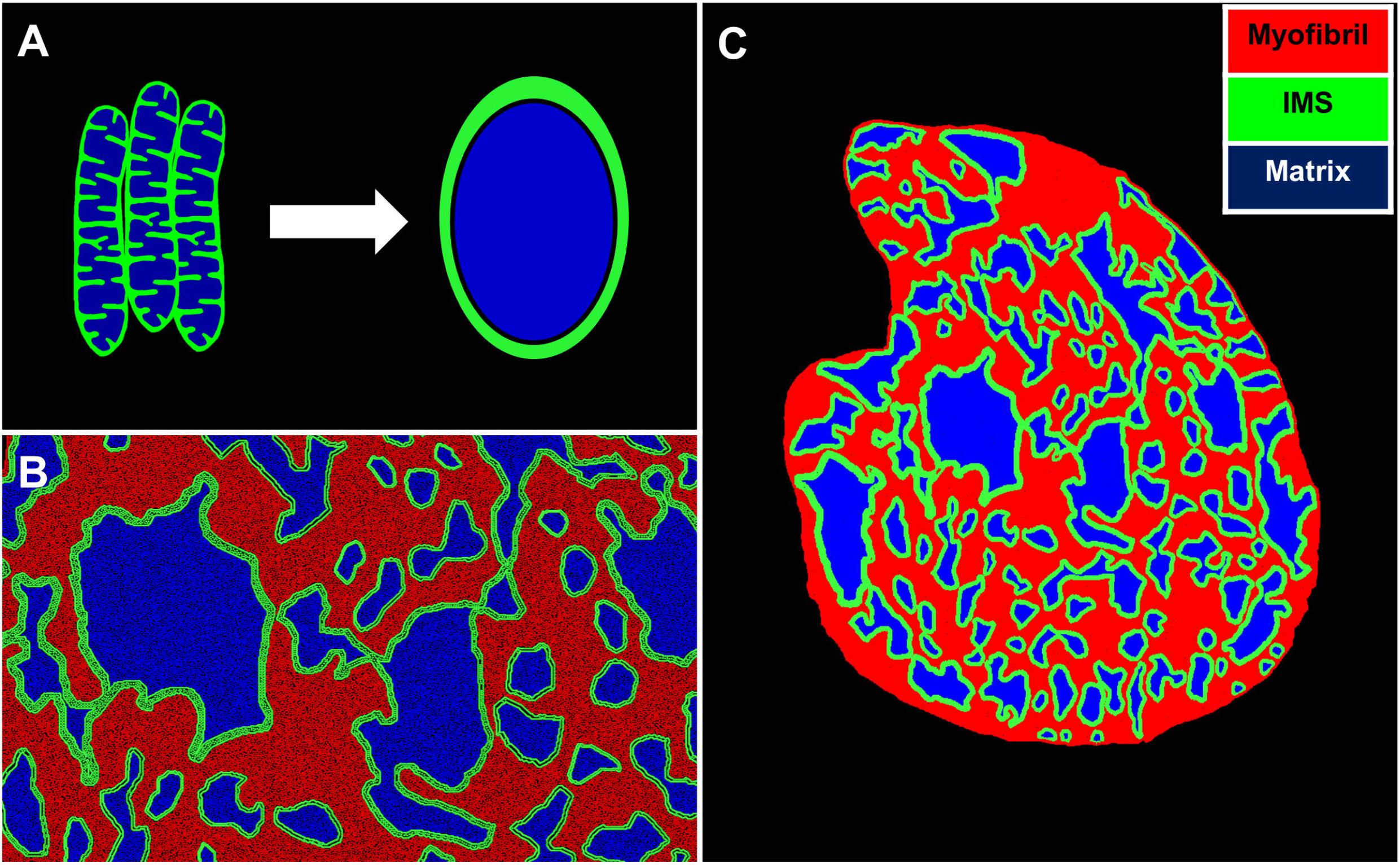
Model assumptions and FE mesh generation from EM image cross-sections. (A): We assumed that individual mitochondria that cluster together will share a common inner membrane space. (B): Based on this assumption, each mitochondrial cluster was divided into two regions: an IMS region, depicted in green and a matrix region, depicted in blue. A third region depicted in red covers the myofibrils present in the cell. (C): Image of a FE mesh developed from the EM image of cross-section 1 with the three different regions.

Assuming there is metabolic connectivity between juxtaposed mitochondria, we divided every mitochondrial cluster present in a cross-section EM image into two regions: (i) an IMS region where we implemented partial differential equations (PDE) to simulate diffusion of phosphagens; (ii) and a mitochondrial matrix region with negligible metabolite diffusivity (Fig 2B). The mitochondrial matrix reactions associated with oxidative phosphorylation (OXPHOS) were simulated using an ordinary differential equation (ODE) model developed by Beard [18]. These ODEs were solved at each computation node on IMS regions and coupled with PDEs for metabolite diffusion within the IMS region. As such, we developed a FE mesh from EM images of section 2 using this protocol (Figure 3C).

**Fig 3.**
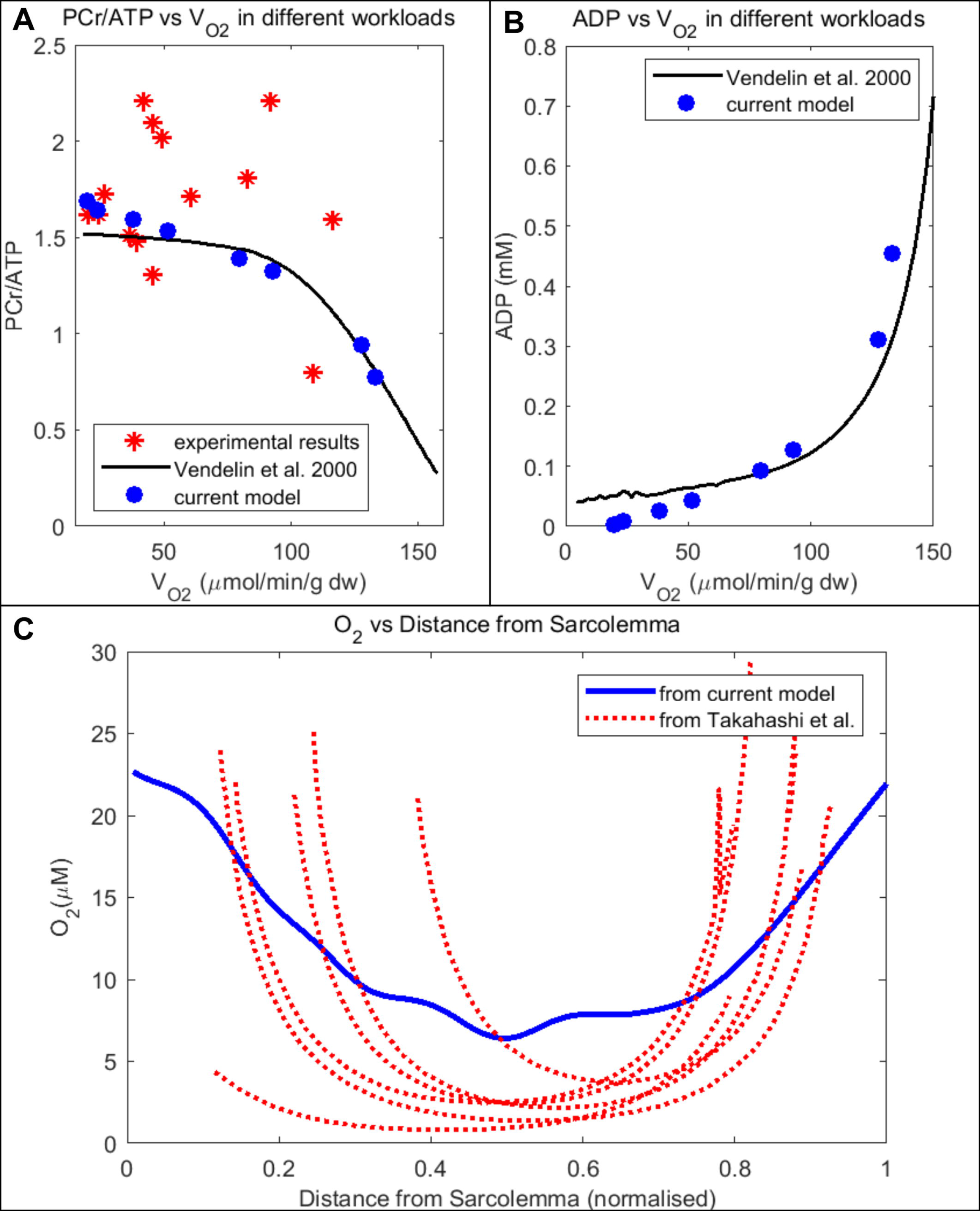
Model validation with experimental results. (A): Comparison between our model-predicted, spatially averaged ATP/PCr ratio vs V_O2_ with experimental results from Valdur Saks et al. [24] and model predictions from Vendelin et al. [10] (B): Comparison between our model predictions for spatially averaged ADP vs V_O2_ with predictions from Vendelin et al.’s [10] zero dimensional model (C): Comparison between model predicted radial profiles of O_2_ with intracellular PO_2_ levels calculated by Takahashi et al. [26] in six different isolated cells.

We assumed a uniform distribution of various enzymes like cytosolic creatine kinase enzymes, adenylate kinase and contractile proteins over the myofibril region within the FE mesh. The equations describing ATP consumption in the myofibril was modelled as a Michaelis–Menten function previously developed by Wu et al [19].

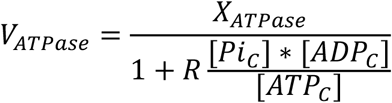

Here, [*Pi_C_*] denotes the cytosolic concentration of inorganic phosphate, *R* is a constant mass-action ratio and X_ATPase_ is a model parameter that can be varied to simulate steady state ATP consumption at various workloads. In order to simulate the action of enzymes such as adenylate kinase and various cytosolic (M-CK) and mitochondrial (mt-CK) creatine kinases, the model also incorporated equations describing the enzyme kinetics from a reaction-diffusion model of cardiac energy transfer described in [10].

We modelled free diffusion of Pi, PCr and Cr between the shared IMS region and the myofibrillar region in the segmented images. On the other hand, diffusivity of ATP, ADP and AMP were set to a value 1% of that in the IMS region, compared to their cytosolic values. This was done to simulate low permeability of the VDACs (voltage depended anion channels) to these large anions as suggested by previous studies [10, 20, 21]. We also assumed that none of these large molecules can diffuse through the mitochondrial matrix due to intramatrix barriers like cristae structure and protein complexes [22].

Owing to its small molecular size, O_2_ was assumed to diffuse through the mitochondrial matrix with similar diffusivity as in the myofibrils and IMS. Therefore, O_2_ was modelled as a species diffusing equally through all the three regions of the EM images. We imposed a constant Dirchlet boundary condition of O_2_ in the cell membrane to simulate the O_2_ flow from a capillary into the cell.

We further utilized an energy metabolites sensitive model of cross-bridge cycling previously developed by Tran et al [23] to predict the force generation at different points in the myofibril area based on the concentration of phosphagens like ATP, ADP and Pi. The overall model was solved using the finite element method. A detailed description of the model formulation and choice of various parameters and initial/boundary conditions can be found in Supplementary Material (S1 Text).

Before making any biological predictions based on the model, it was critical to validate our model with experimental results. The PCr/ATP ratio is one of the most common experimental parameters measured in NMR experiments on in vivo heart. In these experiments [24], PCr/ATP levels have been reported to remain constant over low and moderate workloads, but declines significantly at high workloads (Fig 3A). As evident from Fig 3A, our model was able to reproduce similar behaviour of PCr/ATP ratio for different levels of O_2_ consumption. In the same Fig 3A, we have also presented a comparison of the model predicted steady state PCr/ATP ratio with similar prediction from the compartmental model of rat cardiomyocyte energetics by Vendelin et al. [10]. Our model results were consistent with the compartmental model results.

Our model also predicted a parabolic relationship between average cytosolic ADP concentration vs V_O2_, similar to the prediction from Vendelin et al.’s computational study (Fig 3B). These results indicate accurate modelling of the mechanism of substrate feedback between the mitochondria and myofibrils. The V_O2_ value of 100 μmol/min/g tissue in the plot corresponded to a high workload level of X_ATPase_ = 0.05 μM/sec. According to Vendelin et al.’s study [10], this V_O2_ value also corresponds to around 60% of the maximal filling rate of isolated rat heart perfused according to the “working heart” protocol of Neely [25]. Therefore, we have used X_ATPase_ = 0.05 μM/sec to simulate high physiological levels of workload in all the subsequent simulations.

In order to make accurate predictions about spatial variation in metabolite concentrations or reaction rates, the model needs additional validation with experimental results, where spatial distributions of diffusing species in the cells have been measured using live imaging techniques (i.e. fluorescence microscopy). In a previous study, Takahashi et al [26] visualised intracellular gradients of myoglobin oxygen saturation using high spatial resolution spectrophotometry and further converted the observed oxygenation level into partial pressure of O_2_ (PO_2_) using linear regression of data and the Hill equation. Fig 3C represents a comparison between Takahashi et al’s data and model estimations for radial profile of PO_2_ inside an isolated rat cardiomyocyte. In the experiment, individual cells were subjected to a superfusing O_2_ pressure of 15.2 mmHg under a high O_2_ consumption rate. We simulated high O_2_ consumption in a similar range in cross section 1. Subsequently, the model predicted image of PO_2_ profile was reduced in resolution and convolved with a confocal microscope point-spread-function to generate a simulated confocal image (S2 Fig), similar to a previous study [27]. Figure 3C shows that the concentration profile in the simulated confocal image of O_2_ distribution follows a similar profile to the radial PO_2_ gradient experimentally measured by Takahashi et al. [26]. Both the model and experimental measurements demonstrate steep concentration gradients proximal to the sarcolemma and shallow gradients within the cell interior.

The overall analysis presented in Figs 3A-C illustrates that our model successfully reproduces both the whole-cell averaged and spatially profiled experimental results described in literature.

### Heterogeneous mitochondrial distribution leads to large gradients in metabolite concentration across the cross-section of the cell

Following model validation with the simulation results from cross-section 1, we used the same protocol to simulate cardiac energy metabolism in the three cross sections presented in Figs 1B-D under a high workload (V_O2_ = 100 μmol/min/g tissue). It was assumed that the cell is exposed to a normal oxygen pressure from capillaries, and the concentration of O_2_ at the sarcolemma was maintained at 45 μM based on Mik et al.’s experimental study [28].

Profiles of cytosolic ADP/ATP ratio, Pi and ATP hydrolysis rate (JATPase), corresponding to these cross sections (Fig 4) show significantly large gradients (600-1000 μM) of Pi concentration across the cell. On the other hand, ATP and ADP show negligible spatial gradients, in the order of 10 μM. The ATP hydrolysis rate, being a function of both Pi and ADP/ATP, varies by a margin larger than the ADP/ATP ratio variation but lower than the gradients of Pi. By comparing the different cross-sections (Fig 4), we also observe that section 1 has a higher average level of ADP and Pi compared to the other two sections for the same cytosolic X_ATPase_ activity. Section 2 and 3 also show a larger gradient in the V_ATPase_. These differences can be explained by the lower mitochondrial area fraction of section 1 and smaller spatial variation in mitochondria density distribution as shown in fig 1E. Although section 1 is closer to section 2 than section 3 (12.5 μm vs 25 μm of longitudinal distance), section 1 is located near a branching point of the cell unlike other two sections. Besides similar average values, cross-sections 2 and 3 also have very similar spatial profiles of Pi. The Pi concentration is higher in the upper corner of these cross sections and it gradually increases towards the lower left of the sections. These results are consistent with our initial assumption of symmetric metabolic landscape between different cross-sections of the cell.

**Fig 4.**
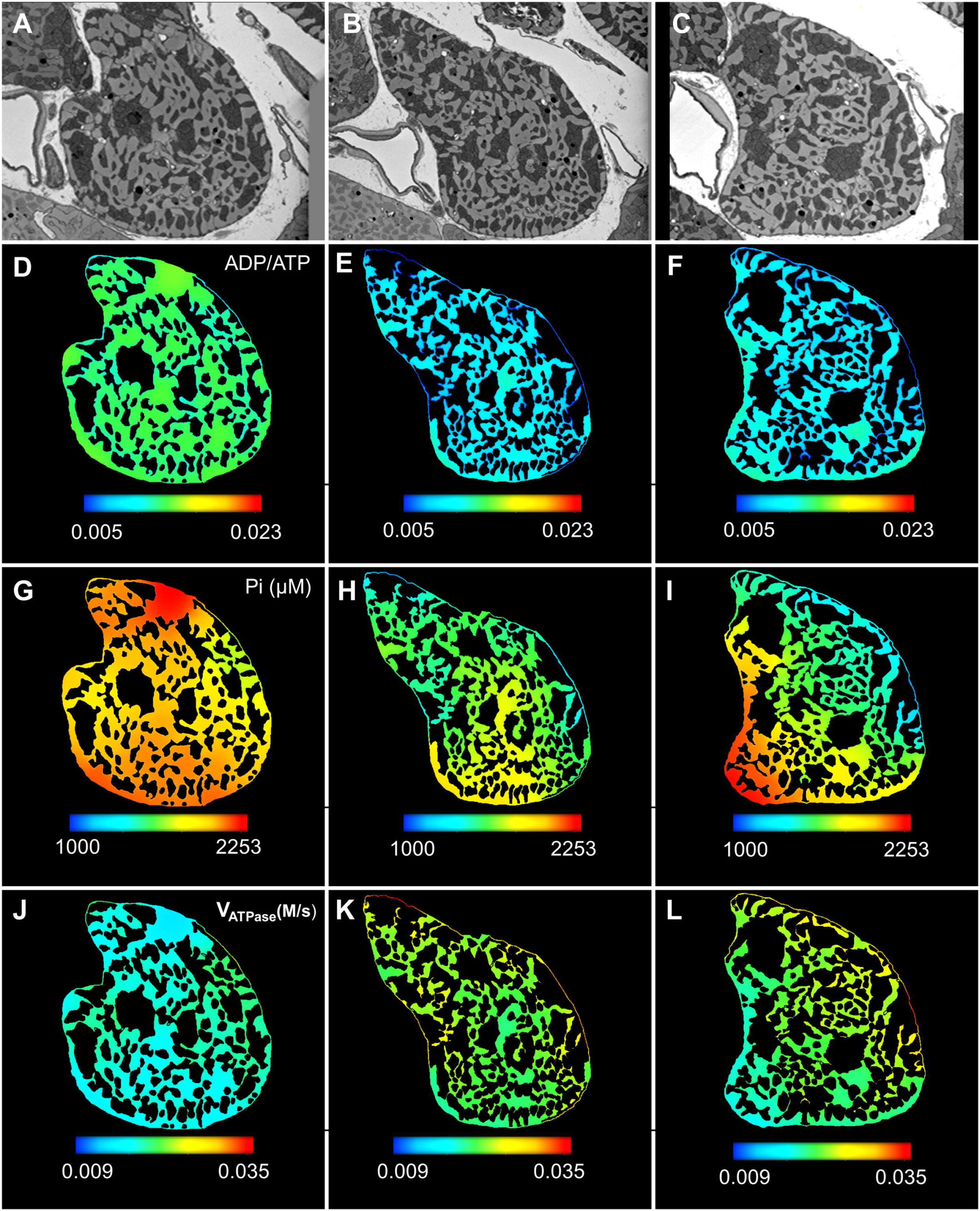
Spatial variation in model predicted cytosolic phosphagen levels and reaction rates represented using colour spectrum. All the figures were generated by using the same range of colour spectrum for a particular phosphagen. (A-C): EM images of sections 1-3 (D-E): Distribution of ADP and ATP in the cross-sections represented using the ADP/ATP ratio. The ADP/ATP ratio remains uniform throughout the cross-sections (G-I): Inorganic phosphate (Pi) shows large spatial variation at the order of 1 mM. (J-L): Spatial variation in the ATP hydrolysis rate in the myofibrils corresponding to X_ATPase_ = 0.05 μM/sec.

In the Fig 5A we have presented a boxplot of the model predicted gradients of all the diffusing metabolite levels present in cross section 1. The differences in spatial gradients of phosphagens can be attributed to the activity of creatine kinase (CK) enzyme isoforms in the IMS and myofibrils. Based on previous work by Vendelin et al [10, 21], we assumed ADP, ATP and AMP diffusivity in the IMS region is 1% of their corresponding diffusivity in the cytosolic space. Despite restricted diffusivity, the estimated ADP and ATP gradients were less than one tenth of the gradients present in PCr and Cr (Fig 5A). These results point towards the significance of the cytosolic MCK enzyme reaction, which acts as a spatial buffer of the ADP/ATP ratio by converting excess ADP into ATP at the cost of the PCr reserve of the cell. We contrasted the PCr and Cr concentration at different regions of cross section 1 with the corresponding local mitochondrial density (Fig 5B). This model predicts that myofibril regions with lower mitochondrial densities contain a lower reserve of PCr. The model predictions are also consistent with the accepted role of the PCr shuttle, in facilitating a rapid metabolic exchange of the equivalent energy of ATP between the IMS and myofibrils.

**Fig 5.**
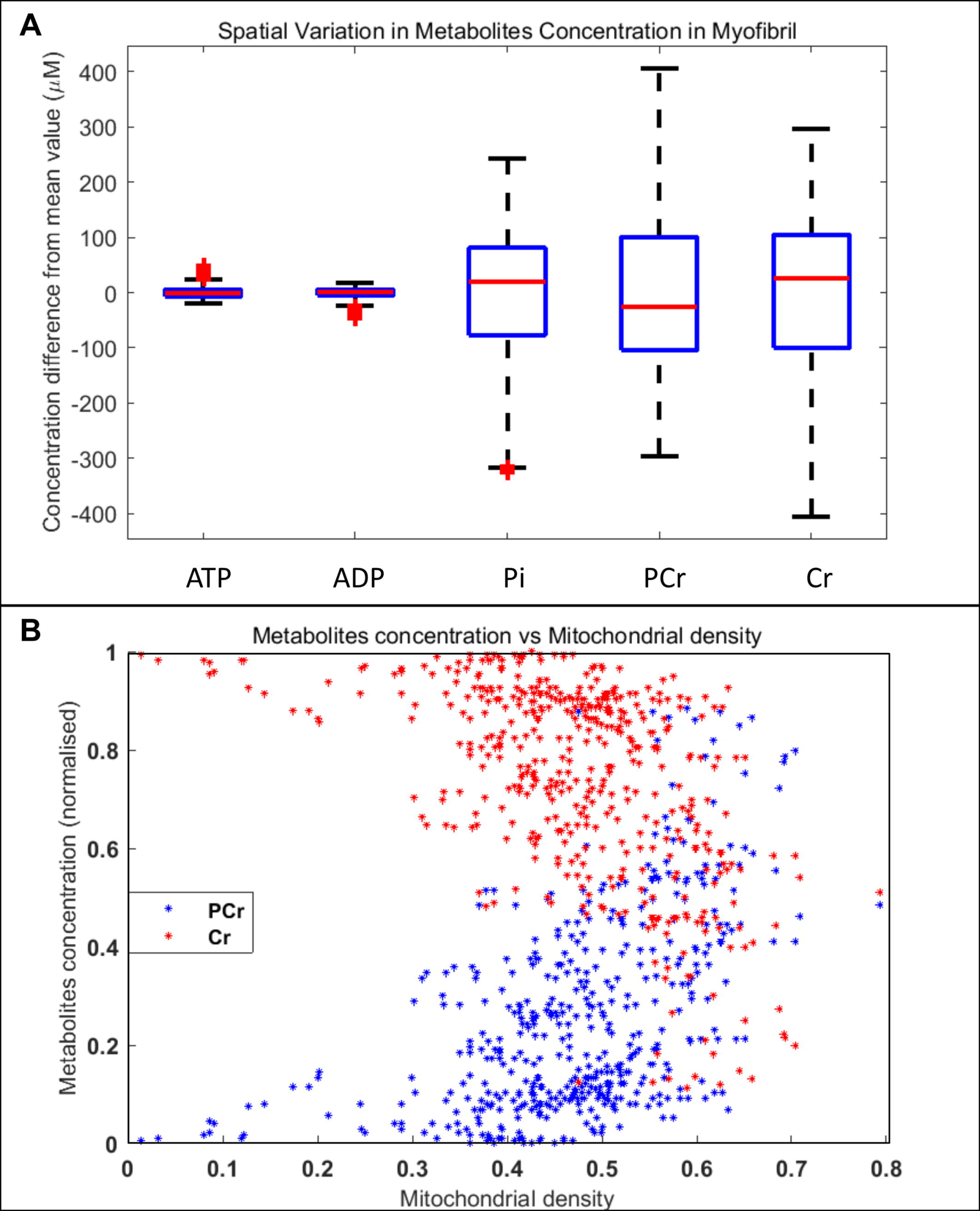
Spatial variation in phosphagens concentration in cross section 1. (A): Phosphagen concentrations were calculated at all the myofibril points in the mesh and subtracted from the spatially averaged concentration of each phosphagen (Phosg_ave_). Box plots represent the statistical distribution of the difference between phosphagen concentration at each point and Phosg_ave_ value of each phosphagen. The zero value corresponds to the Phosg_ave_. (B): Normalized concentration of PCr and Cr displayed as a function of local mitochondrial density at various points in the cell. PCr and Cr concentrations were normalized by a factor equivalent to the difference of PCr and Cr maximum and minimum concentrations in cross section 1.

### Force dynamics is insensitive to heterogeneity in concentration of phosphagens and distribution of mitochondria

To analyse the impact of the spatial distribution of energy phosphagens on mechanical force production, we have used Tran et al.’s [23] cross-bridge cycling model to simulate isometric twitches at each of the discretised nodes. The isometric twitches are activated by a prescribed Ca2+ transient. It was assumed that the average value of various metabolite concentrations remains unchanged during a twitch, and accordingly the behaviour of the cross section was modelled as a function of cytosolic ATP, ADP, Pi, MgATP, MgADP and pH. The peak twitch force (F_peak_) was simulated (Figs 6A-C) for the cross sections, as were twitch durations (Figs 6D-F), defined as the duration that the force was above 5% of F_peak_ (t_95_). F_peak_ shows slight spatial variation in sections 2 and 3 (~7% difference between maximum and minimum spatial values), while it is almost uniform across section 1 (<3% difference). Similarly, differences between the maximum and minimum values of t_95_ is around 12% in section 2 and 3, while it is around 5% in cross-section 1. These results imply that the force production is relatively insensitive to the large gradients in inorganic phosphate observed in Figs 4 and 5. The results also predict that rapid diffusion of PCr and Cr, coupled with CK enzyme reactions in myofibrils and IMS, is sufficient to maintain a uniform force generation across the cell cross section despite the heterogeneous distribution of mitochondria.

**Fig 6.**
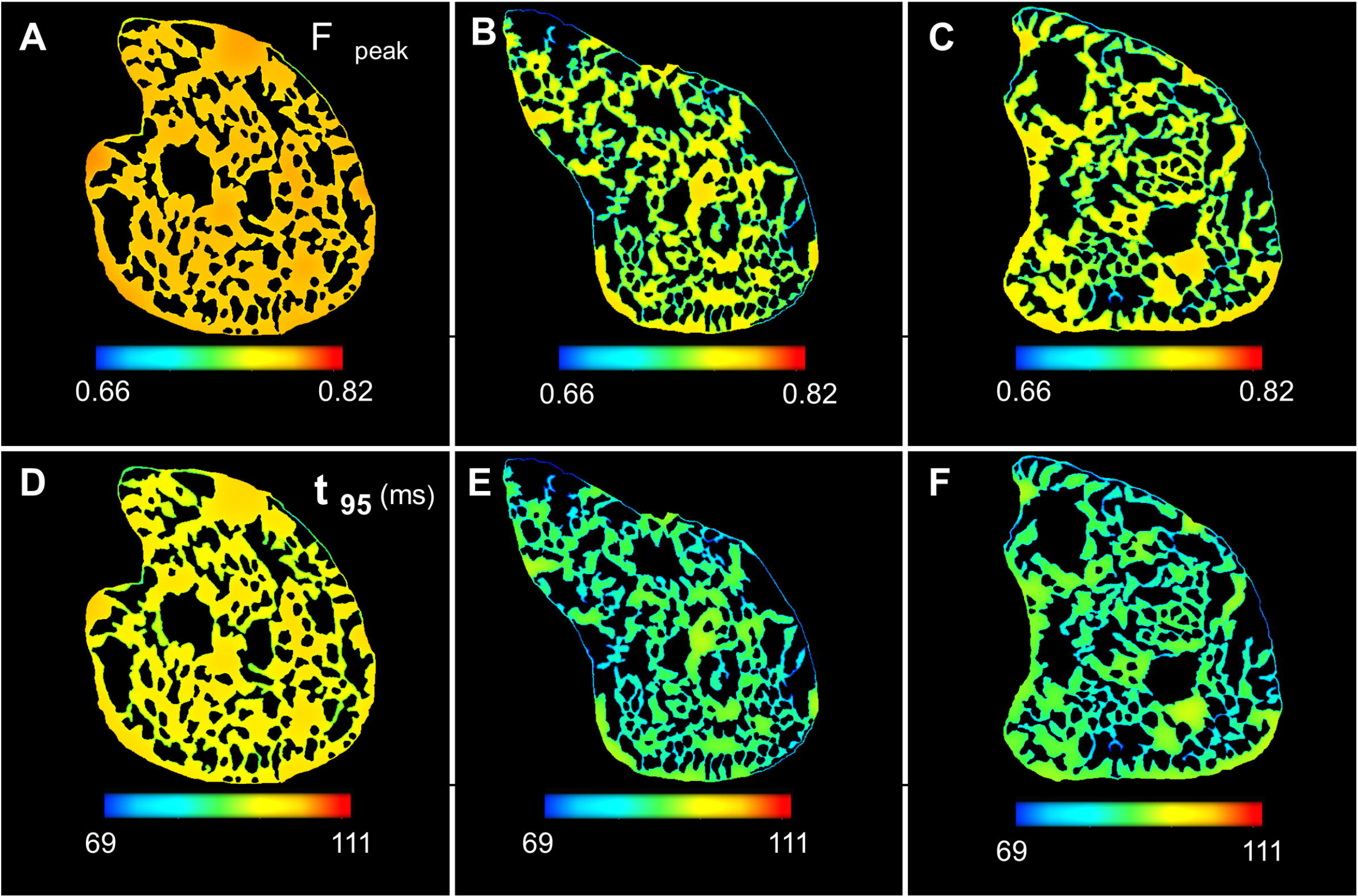
Force dynamics of the cell corresponding to the model predicted phosphagens distribution. (A-C): F_peak_ is uniform across section 1 and exhibits slight spatial variation in section 2 and 3. (D-E): t_95_ shows similar spatial gradients across the cross sections.

### Diffusion of PCr & Cr is not sufficient to maintain uniform force dynamics under hypoxic conditions

According to previous studies [29, 30] on isolated rat cardiomyocytes, cardiac energetics is impacted when the extracellular O_2_ diminishes below a margin of 20 μM, as the mitochondrial oxidative phosphorylation system is limited. In an in-vivo non-invasive study on wistar rat hearts [28], Mik et al. found an average PO_2_ of 35 mm Hg (roughly equivalent to 45 μM concentration assuming an O_2_ solubility coefficient 1.35×10−6 M/mmHg [31]) present over the surfaces of cardiac mitochondria, which is well above this margin. However, Mik et al. also found that a significant percentage (25%) of in-vivo mitochondria was exposed to PO_2_ in the range of 10−20 mm Hg under normal O_2_ supply. Reducing the O_2_ supply to the heart resulted in an even larger fraction of cardiac mitochondria (40%) with low surface PO_2_ in the range of 0-10 mm Hg. Mik et al.’s findings were consistent with predictions from previous modelling studies by Beard et al. [31] and Groebe et al. [32]. Both of these modelling studies predicted that cardiac myocytes in-vivo can be exposed to a variety of PO_2_ depending on location of the cells relative to capillary networks.

Therefore, we investigated the effect of low oxygen on phosphagens distribution in the 2D cross-sections. As shown in the previous results, we first applied a Dirichlet boundary condition with O_2_ concentration of 45 μM along the cell membrane, which represents the majority of cells present in heart. In the second simulation we imposed a 15 μM oxygen concentration along the cell membrane, representing the cells exposed to lower levels of oxygen supply (hypoxic condition). Both the simulations corresponded to the same high workload levels.

Profiles of O_2_ concentration in the cardiomyocyte, corresponding to these two simulations, are shown in Fig 7A. In both the cases, the O_2_ distributions show steep gradients from the sarcolemma. While in the first case O_2_ concentration remains above 20 μM throughout the cell, our model predicts an anoxic cell core under hypoxic conditions. This prediction is consistent with observations of Takahashi et al. [26, 33] on isolated cardiomyocytes as well as Mik et al.’s study [28] on mitochondria in the in-vivo heart. To better understand the effects of an oxygen deprived cell core on mitochondrial electron transfer system (ETS), we have plotted the mitochondrial complex IV reaction rate (V_C4_) as a function of the IMS space (Fig 7B). Complex IV is the main site of oxygen consumption in the ETS. As expected in normoxia, activity of V_C4_ is nearly uniform across the cell cross section, with small variations due to mitochondrial cluster sizes. However, in the hypoxic simulation, a negligible amount of O_2_ passes to the interior of the cell, which in turn inhibits the J_C4_.

**Fig 7.**
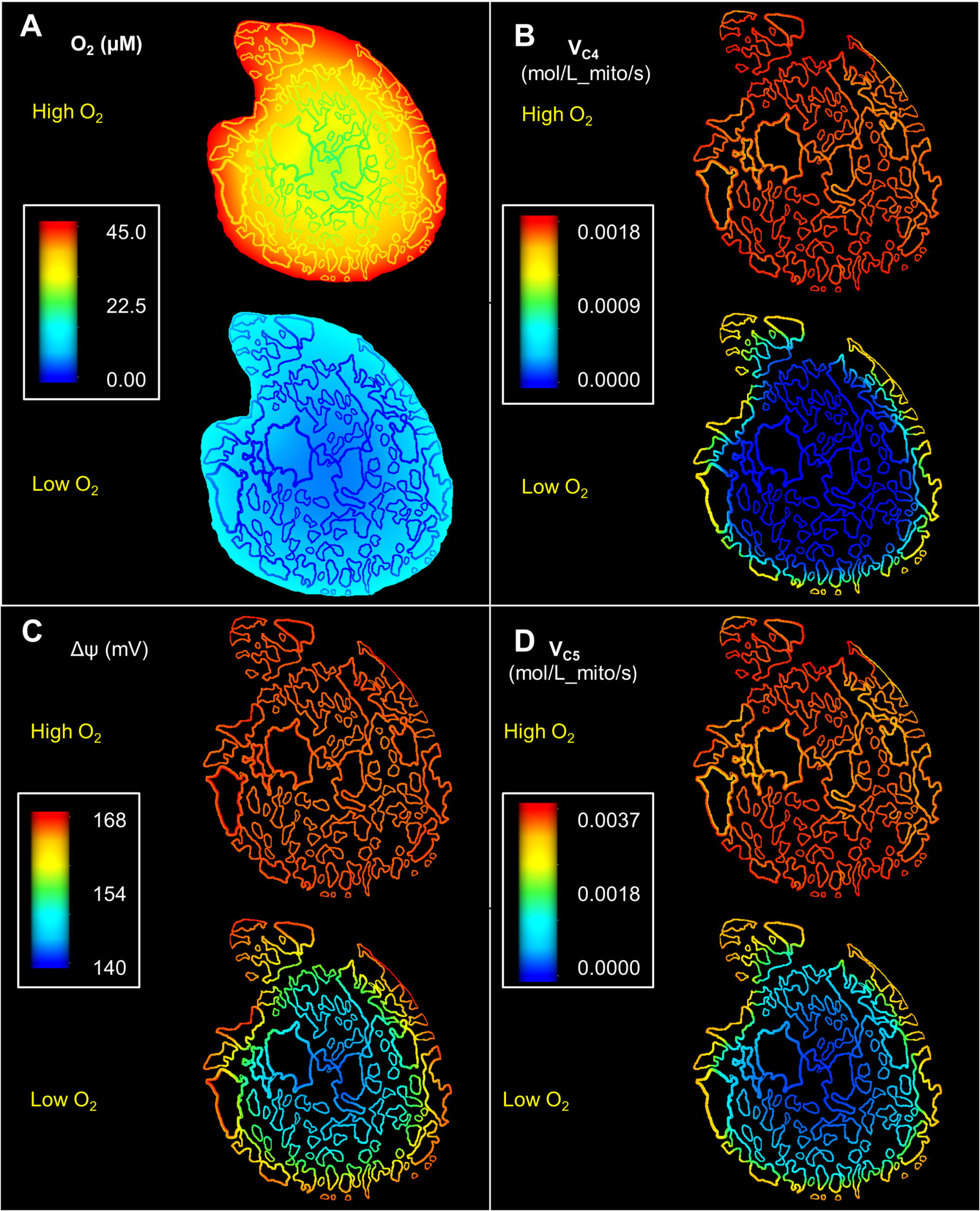
Spatial variation in model predicted mitochondrial reaction rates and O_2_ profiles for cross section 1. All the figures were generated by using same colour spectrum range. (A): Steady state O_2_ profiles showing steep O_2_ gradients in the vicinity of the sarcolemma. In the hypoxic case, it also leads to the formation of an anoxic core with negligible O_2_ level (B): The reduction of oxygen at complex IV is inhibited in hypoxia, while it is uniform across the section in normoxia. (C): Hypoxia leads to a steep gradient in the model predictions for mitochondrial membrane potential (D): Under normoxia Complex V (F_1_-F_0_ ATP synthase) reaction rate is nearly uniform across the cell with slight variation near the larger mitochondrial clusters. However, ATP synthesis at Complex V is inhibited in the anoxic core under hypoxia.

We also investigated the influence of intracellular O_2_ profiles on mitochondrial membrane potential (Δψ) for both the normoxic and hypoxic simulations (Fig 7C). In the first case, Δψ was nearly uniform across the cell. This uniform distribution of Δψ is clearly in accordance with the uniform profile of V_C4_ observed in Fig 7B. However, in the second case, Δψ is significantly depressed in the core of the cell. This spatial depression of Δψ can be attributed to the decreased ETS flux in the centrally located IMF mitochondria. The lower row in Fig 7D demonstrates the effect of this depression in Δψ on reaction flux of complex V. The rate of ATP synthesis at complex V follows the same spatial profiles as Δψ. As a result, ATP synthesis declines in the centre of the cell in hypoxia.

The diminished rate of ATP synthesis in the mitochondria at the core of the cell also had a significant impact on transportation of ATP/ADP though the phosphocreatine shuttle (Fig 8). For instance, the reaction rate of mtCK in the forward direction (ATP + Cr → PCr + ADP) was around 15% lower in the central parts of the cell compared to the periphery under hypoxia (Fig 8A). As a result, the model predicts a strong radial gradient in the ADP/ATP ratio in the hypoxic simulations. This is in contrast to the simulations of normoxic situation where both the mtCK reaction rates and the ADP/ATP ratio shows negligible spatial variation (Fig 3A and 9A). Fig 9B similarly demonstrates a radial profile in the ATP hydrolysis rate for the second case. The abundance of ADP in the anoxic cell core also leads to an elevated cytosolic MCK reverse reaction rate (PCr + ADP → ATP + Cr) which is substantially higher in the anoxic core compared to the periphery of the cell (Fig 8B). Together these results suggest that activity of mtCK and MCK is not sufficient for maintaining uniform ADP/ATP under limited O_2_ supply.

**Fig 8.**
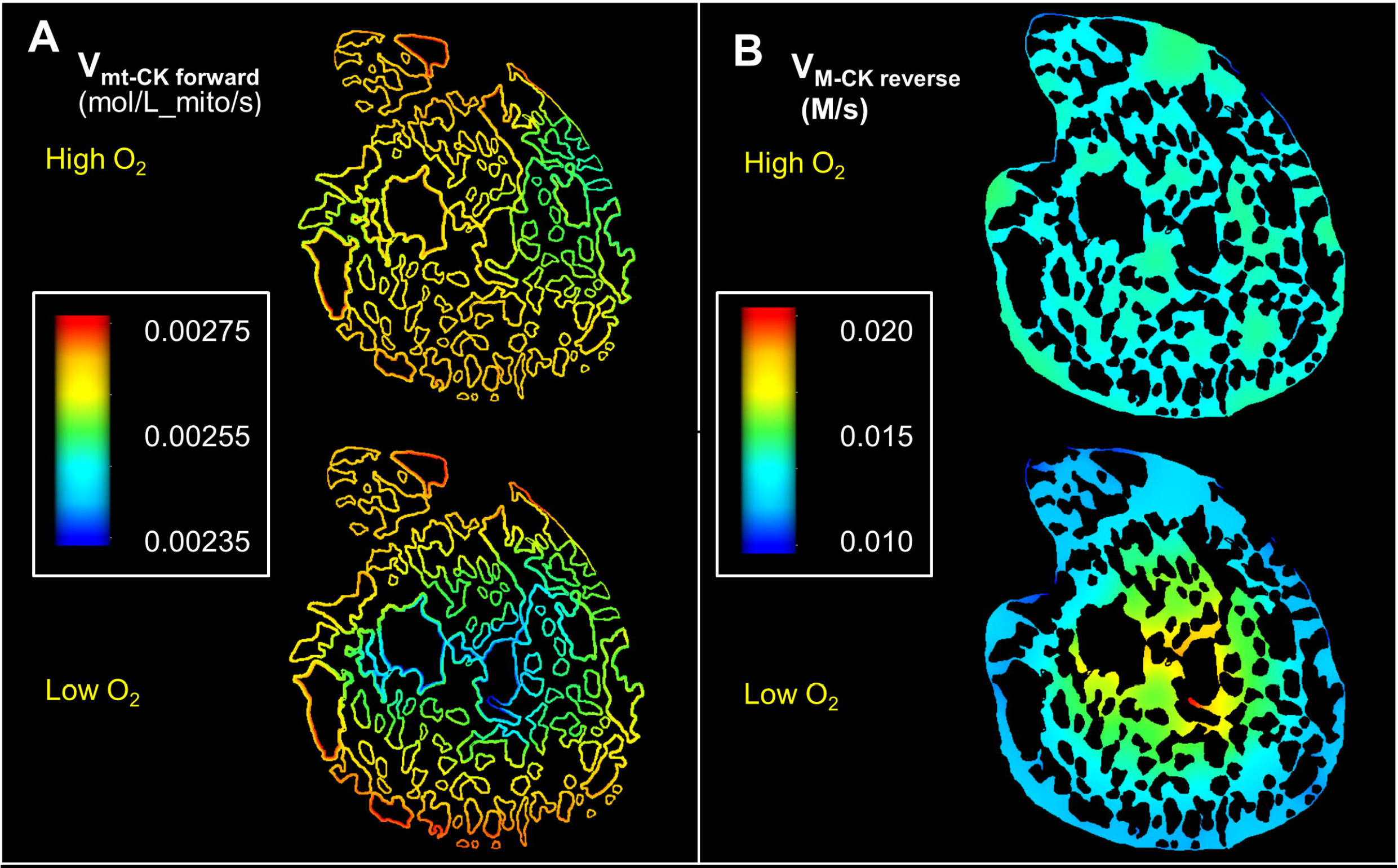
Spatial variation in rates of CK enzymatic reactions involved in phosphocreatine shuttle under normoxia and hypoxia. All the figures were generated by using same colour spectrum range. (A): Steady state profiles of reaction rates of mitochondrial mtCK in forward direction (ATP + Cr → PCr + ADP) (B): Steady state profiles showing reaction rates of myofibrilar MCK in reverse direction (PCr + ADP → ATP + Cr)

**Fig 9.**
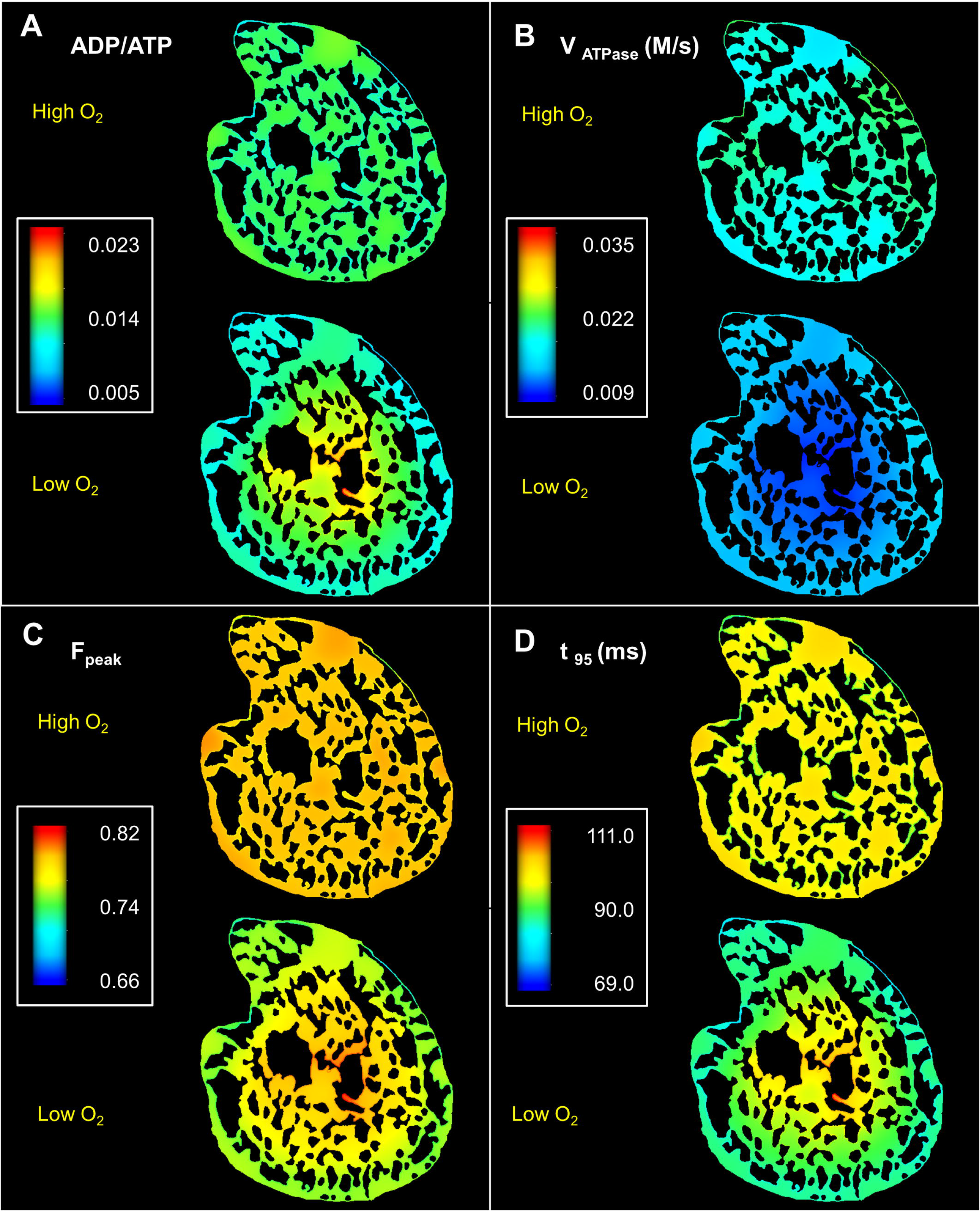
Spatial variation in cytosolic metabolite concertation and force dynamics under normoxia and hypoxia. (A): Steady state profiles of ADP/ATP ratio. ADP/ATP is significantly higher in the anoxic cell core under low O_2_ supply (B): ATP hydrolysis rate is similarly lower in the cell core under hypoxic condition (C): Peak twitch force (F_peak_) is significantly higher in the cell core under hypoxia (D): Increased F_peak_ is complemented by increased twitch duration in the cell core, which can lead to loss of energy in the form of shear strain.

For the second simulation, F_peak_ is also higher in the anoxic core of the cell and the twitch durations are significantly longer. The difference between the maximum and minimum spatial values of t_95_ is around 23% in the second simulation compared to only 5% in the first simulation. This is a result of lower ATP, and higher ADP and Pi in the centre of the cell - which slows down the cycling rate, but at the same time, the cross bridges spend more time in the force-producing states (due to lower ATP), leading to higher force development. A consequence of having longer twitch durations is that at high stimulus frequencies, the twitch does not have time to completely relax resulting in a higher diastolic force. Thus according to the model predictions, different regions of the same cell cross section will be experiencing different levels of force in both systole and diastole. This may lead to shear strain inside the cell and wastage of metabolic energy through production of heat – which might not be a favourable situation during a high cardiac workload.

These results imply that unlike under normoxic conditions, diffusion of Cr and PCr and associated CK enzyme activities is not sufficient to maintain a uniform ATP/ADP concentration and force dynamics under hypoxic conditions.

## Discussion

We have developed a computational model of cardiac bioenergetics based on EM images to study the impacts of the heterogeneous distribution of mitochondria on the energy metabolic landscape of the cell. Note that this differs markedly from the earlier held concept that cardiomyocyte mitochondria are arranged within rigid crystalline lattice networks, which is thought to enhance diffusion and exchange of compounds such as the phosphagens. When less structured mitochondrial arrangements are incorporated into models, our results reveal that significant gradients in the phosphagen concentrations are likely. While concentration gradients are currently difficult to observe directly in experimental studies, previous computational studies, defined over idealised crystal lattice-like geometries, predicted smaller concentration gradients between adjacent mitochondria and myofibril units. Our current study indicates that significant concentration gradients may occur over the entire cross-sectional area of the cell, that this is dependent on localised mitochondrial densities at a given point in the cell, and that these gradients appear acutely sensitive to hypoxia, such as would occur in infarcts.

Our predictions are consistent with the theoretical role of creatine kinase (CK) in maintaining ATP homeostasis across the cell. Cytosolic ATP pool not only act as a source of ATP for the cross-bridge cycle, but it also supports ion channel activities (e.g. SERCA pumps) and can act as divalent metal ion chelator [34], crucial to the viability of the cell. According to a previous modelling work [35] SERCA pumps, which are intrinsic to cellular Ca^2+^ management and EC-coupling, appear to be highly sensitive to the ATP/ADP ratio in the cell. Therefore, it is vital for cardiomyocytes to maintain a uniform ATP and ADP concentration, in spite of the heterogeneous distribution of mitochondria.

We used an O_2_ diffusivity value (D_O2_) of 2.41×10^−5^ cm^2^/s in our current simulation based on previous modelling work of Beard et al. [31]. This diffusivity value corresponds to the overall oxygen diffusion through mammalian striated muscles [36], and can be considered as an average diffusivity through all the organelles present inside a cardiomyocyte. Therefore, we did not separately model O_2_ transport through the transverse tubules which are near uniformly distributed along the Z disks [37]. We also neglected the effect of myoglobin facilitated oxygen diffusion since diffusivity of myoglobin is reported to be only a small fraction (<1/10_th_) of the free O_2_ diffusivity in cardiomyocytes [38]. Despite these assumptions, our model prediction of radial profiles in oxygen is consistent with previous experimental and theoretical works [26, 32]. Our model indicates that in a normoxic state, profiles of O_2_ have no effect on the metabolism, and that the phosphagen distribution is dictated by the mitochondrial distribution.

However, in hypoxia, our model predicts a significant impact of O_2_ profile on the phosphagen distributions and the effect of the heterogeneous mitochondrial distribution is minimal. The level of mitochondrial membrane potential in the core of the cell was substantially lower than the periphery of the cell. The ATP/ADP in the cell core was similarly depressed due to a lack of O_2_. However, these predictions are inconsistent with findings from Takahashi et al.’s 2008 work [33] exploring isolated cardiomyocytes. Takahashi et al. showed that isolated cells sustain depleted, but near homogenous levels of ATP and Δψ, in spite of formation of an anoxic core, when the respiration rate is elevated under a hypoxic condition. Takahashi et al. also suggested that CK mediated diffusion of PCr and Cr from the sub-sarcolemmal space supports ATP supply and Δψ levels in the anoxic core for a substantial time interval before necrotic cell death. Our results contrast with this view, as they indicate that CK reaction and diffusion of PCr and Cr is not sufficient to generate a homogenous metabolic landscape under hypoxia. This inconsistency between our model predictions and experimental data suggest that other mechanisms may be at play to ensure uniform ATP supply. Other mechanisms could include glycolysis, creatine phosphate stores or Δψ conductivity between sub-sarcolemmal and inter-myofibrilar mitochondria. Glancy et al recently proposed that Δψ can be transmitted across the cell through mitochondrial networks in both skeletal [39] and cardiac muscle cells [8]. Transmission of Δψ might be facilitated through IMJs present between adjacent mitochondria. Glancy et al. also proposed that membrane potential conduction via mitochondrial reticulae is a dominant pathway for skeletal muscle energy distribution.

In the current study, we assumed that there is no conduction of membrane potential between adjacent mitochondria, and accordingly modelled Δψ as a spatially distributed non-diffusing variable without any direct interaction between adjacent mitochondrial points. With this modelling approach, we did not find any significant difference in the mitochondrial Δψ values across the cross section of the cell in normoxia. These results are consistent with Glancy et al.’s [8] observation that mitochondrial Δψ conductivity is much weaker along the cell cross section compared to the longitudinal axis of the cardiomyocytes. We propose that, in normoxia, the rapid diffusion of cytosolic PCr and Cr is sufficient to maintain a uniform Δψ as well as myofibrillar ATP distribution across the cell cross section without the need for Δψ conduction. However, our results also imply that in a hypoxic cardiomyocyte, the phosphocreatine shuttle is not sufficient to maintain a uniform distribution of ATP. Δψ conduction might play a role in maintaining uniform levels of intracellular ATP across the cell cross-section. In the future works, it need to be investigated whether hypoxia can lead to formation of larger number of IMJs along the cell cross section which can subsequently lead to higher Δψ conductivity compared to normoxic condition.

The total cellular PCr pool and activities of mitochondrial CK enzyme are severely impacted in a number of metabolic diseases, such as diabetic cardiomyopathy. According to previous studies [40–42], mtCK and MCK activities are suppressed by a substantial margin in diabetes (35%-50%). Without normal CK activities, homogenous ATP distribution within cardiomyocytes can be severely impacted due to heterogeneous mitochondrial distribution. Our previous studies also revealed that mitochondrial arrangement in type 1 diabetes is significantly irregular than control cells due to mitochondrial fission and clustering [6], which can also affect the distribution of cellular phosphagens. In future work, we will study these potential changes in metabolite distribution and their effect on force dynamics using a 3D geometry based FE model of cardiac bioenergetics. Future studies will also require a modification in the currently used model of mitochondrial membrane potential to incorporate the conduction of membrane potential.

## METHODS

### Tissue sample preparation and serial block face imaging

The tissue sample used for SBF-SEM imaging was collected from the left ventricular wall of a sixteen-week-old male Sprague-Dawley rat at the University of Auckland, New Zealand. We followed the guidelines approved by the University of Auckland Animal Ethics Committee (for animal procedures conducted in Auckland, Application Number R826) for the process.

Following euthanasia of the Sprague-Dawley rat, the heart was excised and quickly cannulated and connected to a Langendorff apparatus operating at a hydraulic pressure of 90 cm. We perfused the heart with a Tyrode solution including 20 mM 2,3-Butanedione Monoxime for 2-3 minutes. This was followed by perfusion with 2.5% glutaraldehyde, 2% paraformaldehyde and 50 mM CaCl2 in 0.15 M sodium cacodylate buffer. We dissected a tissue block from the left ventricular free wall and stored in the same fixative that was precooled on ice for 2 hours. The fixative was subsequently replaced with 2% osmium tetroxide and .8% potassium ferrocyanide in 0.15 M sodium cacodylate and left overnight. Afterwards, we stained the block with ice-cold 2% uranyl acetate for 60-120 minutes. Following this, the block was washed of excess uranyl acetate and punched into a 1.5 mm diameter sample. The sample was then progressively dehydrated with ethanol followed by transition to room temperature in acetone. The sample was finally embedded in epoxy resin (Durcupan ACM resin from EM sciences).

We trimmed the tissue sample to a square block face of 1 mm^2^ and 300 μm deep with fibers running perpendicular to the future cutting face. The sample was subsequently mounted on a Teneo VolumeScope (FEI,Hillsborough, USA) and imaged in low vacuum mode (50 mbar) with a 10 nm pixel size. We acquired a total of four thousand and nineteen serial sections of 50 nm in thickness. The sections were then aligned using IMOD. We rotated the sections by 90 degrees around the Y axis and binned the data to an isotropic voxel size of 50 × 50 × 50 nm. A detailed description of the animal procedures, sample preparation and imaging techniques is available at our previous publications [14, 15].

### Model development and computational methods

As discussed earlier, the results section provides a brief description of the model development, while the supplementary document S1Text contains description of all the PDEs, ODEs and algebraic equation used in the model. The finite element implementation of the model was developed using reaction-diffusion module of OpenCMISS [43], an open source software package for FE simulation. Besides PDEs, the OpenCMISS model also contains ODEs and algebraic rate equations that have been described in CellML [44] models – which were solved using the strang operator splitting method. We offer the OPECMISS implementation of the FE model for research use at *https://github.com/CellSMB/cardiac_bioenergetics*.

The FE models were solved on an IBM iDataplex x86 system. Individual FE models contained 300,000 FE nodes on average and required 20 hours of runtime for simulating every 1000 milliseconds of heartbeat. The results in this paper correspond to steady state solutions at 3 seconds of heartbeat time.

## Acknowledgments

The authors would like to acknowledge Mr. Akter Hussain for the help with segmentation of EM images, and Melbourne Bioinformatics for the support in executing the simulations in IBM iDataplex x86 system.

